# DICTrank: The largest reference list of 1318 human drugs ranked by risk of drug-induced cardiotoxicity using FDA labeling

**DOI:** 10.1101/2023.07.06.548029

**Authors:** Yanyan Qu, Ting Li, Zhichao Liu, Dongying Li, Weida Tong

## Abstract

Drug-induced cardiotoxicity (DICT) is one of the leading causes of drug attrition in clinical trials or withdrawal from the market. Many studies have been conducted to detect DICT in the early stage of drug development and clinical diagnosis, but the success is limited, as evident by the high attrition rate at all clinical phases due to DICT. Most of these efforts, if not all, have focused on specific adverse events and/or mechanisms associated with DICT, such as QT prolongation and hERG-related cardiotoxicity. However, given the broad spectrum of cardiotoxicity, it is necessary to develop a reference drug list with a systematic annotation of DICT potential across a large number of drugs and drug classes. Such a list is essential for developing effective DICT diagnostic biomarkers and early prediction strategies using new approach methods (NAMs), including artificial intelligence (AI). By utilizing labeling documents for FDA (U.S. Food and Drug Administration)-approved drugs, we developed a drug classification scheme to prioritize drugs based on their DICT potential. This resulted in DICTrank, which is the largest dataset of drugs annotated with ranked DICT risk in humans. DICTrank categorizes drugs into four categories of DICT concerns by integrating DICT severity and labeling content. The dataset consists of 1318 drugs, classified as follows: Most-DICT-Concern (341), Less-DICT-Concern (528), No-DICT-Concern (343), and Ambiguous-DICT-Concern (106; lacking sufficient information in the labeling document to determine cardiotoxicity potential). DICTrank covers a wide range of drug therapeutic categories. Using this extensive DICT dataset, we discovered that several therapeutic categories were significantly enriched with drugs of Most-DICT-Concern as well as the association of daily dose with drug class. These categories include antineoplastic agents, sex hormones and modulators of the genital system, anti-inflammatory and antirheumatic products, beta-blocking agents, and cardiac therapy. DICTrank represents the largest drug list for DICT to date, and it could contribute to the development of NAMs and AI models for the early identification of DICT risk during drug development and beyond.

## Introduction

Drug development is expensive and time-consuming, for which drug safety has contributed significantly. Approximately one third of all drug discontinuation was caused by severe adverse drug reactions (ADRs), among which drug-induced cardiotoxicity (DICT) is the one of the leading causes of drug failure in the late clinical phase and withdrawal from the market [1,2]. Cardiotoxicity is one of the most common adverse drug effects and it is broadly defined by the National Institutes of Health (NIH) as toxicity that affects the heart. Among 133 drugs withdrawn from the market between 1990 and 2010, 25 cases (18.8%) were caused by cardiotoxicity, following hepatotoxicity (36 cases, 27.1 %) [3].

DICT has been consistently focused on by drug developers, clinicians, and regulators. The critical challenge of DICT management is the broad spectrum of cardiotoxicity patterns, diverse active pharmacological classes, and the complicated interplay between host and drug. DICT is drug-induced injury to the heart, whereas cardiovascular toxicity exhibits a broader scope and encompasses the toxicity affecting the entire cardiovascular system, including the heart, blood vessels, and blood circulation. Major cardiac adverse effects include heart failure, myocarditis, cardiomyopathy, myocardial ischemia, arrhythmias, hypertension, and impairment of cardiac valves [4]. Cardiotoxicity has been observed with a diverse range of drug classes, such as antineoplastic agents, antipsychotics, antidepressants, antibiotics, and neurodegenerative disease agents [5,6]. Drugs in certain therapeutic categories tend to cause more serious DICT than others. For example, anticancer therapies, either chemotherapy or radiation treatment, have been linked to DICT with potential life-threatening consequences [7-10]. As the remarkable progress have been made in cancer therapeutics including new types of anticancer drugs in recent decades, the appropriate management of DICT has gained increasing importance.

To reduce cardiac ADRs, it is crucial to determine DICT’s clinical relevance with therapies, clarify the underlying molecular mechanisms, identify reliable biomarkers, and develop new diagnostic and therapeutic modalities. It is substantially challenging to delineate the molecular mechanisms underlying DICT since cardiotoxicity is particularly broad-spectrum and vary largely among patients [11,12]. In addition to the direct interaction with cardiac electrophysiology, DICT frequently results from disruption of contractility, mitochondrial toxicity, growth factor and cytokine regulation, fibrosis, and thrombosis[13]. Despite the substantial progress in mechanistic study, the underlying mechanisms of DICT have yet to be comprehensively understood, especially in the clinical scenario, which hinders the development of effective approaches to mitigate DICT. Better understanding of DICT mechanisms is important to the development of new drugs and of assays for clinical and preclinical safety testing. However, the lack of a reference drug list of DICT information is the foremost hurdle for mechanistic investigations and development of effective diagnosis and prognosis strategies.

A reference drug list named Drug-Induced QT prolongation Atlas (DIQTA) was reported in 2022, focusing on classification of 428 drugs for QT prolongation risk, one major DICT mechanisms [14]. Cai et al. developed an in-silico approach using combined classifiers for risk assessment of drug-induced cardiovascular complications (not DICT specific), where five reference drug lists were collected separately for hypertension, arrhythmia, heart block, cardiac failure, and myocardial infarction from four publicly available data resources: Comparative Toxicogenomics Database (CTD), SIDER (side effect resource), Offsides (database of drugs effects), and MetaADEDB (adverse drug events database) [15]. Blockage of hERG potassium channel is associated with long QT syndrome. A hERG-related cardiotoxicity dataset composing 2130 compounds was collected by Lee et al., based on which a computational model was built to predict hERG-related cardiotoxicity for drug candidates [5]. Most, if not all, currently existing cardiotoxicity classification are focused on specific adverse events or mechanisms such as hERG-related cardiotoxicity. In both clinical application and drug development, it is necessary to understand a drug’s risk to the heart as a whole instead of individual specific adverse events or mechanisms and to rank the drug by its overall cardiotoxicity potential.

To fill the gap, we developed the Drug-Induced Cardiotoxicity Rank (DICTrank) to classify and rank FDA (U.S. Food and Drug Administration)-approved drugs regarding their potential to cause DICT using FDA drug labeling documents, which have previously been used to classify drug-induced liver injury (DILI) [16-18]. In addition, we elucidated the potential utility of DICTrank in research and clinical application.

## DICTrank Development

The DICT classification scheme proposed in this study was developed based on FDA-approved drug labeling, which is a compilation of information about a drug necessary for its safe and effective use. The drug labeling is written mainly for healthcare practitioners, approved by the FDA, and regulated by law (www.accessdata.fda.gov/scripts/cdrh/cfdocs/cfCFR/CFRSearch.cfm?fr=201.57). Drug labeling provides current drug safety information and is widely recognized as one of the most useful resources to assess drug safety. Generated from clinical trials, post-marketing surveillance, and published literature reports, the information of adverse drug reactions included in drug labeling is most widely utilized by industry, academic and regulatory communities for pharmacovigilance [16,19]. Figure 1 depicts the overall classification approach for evaluating drugs’ potential for DICT, which is further elucidated in the following four sections.

**Figure 1.**
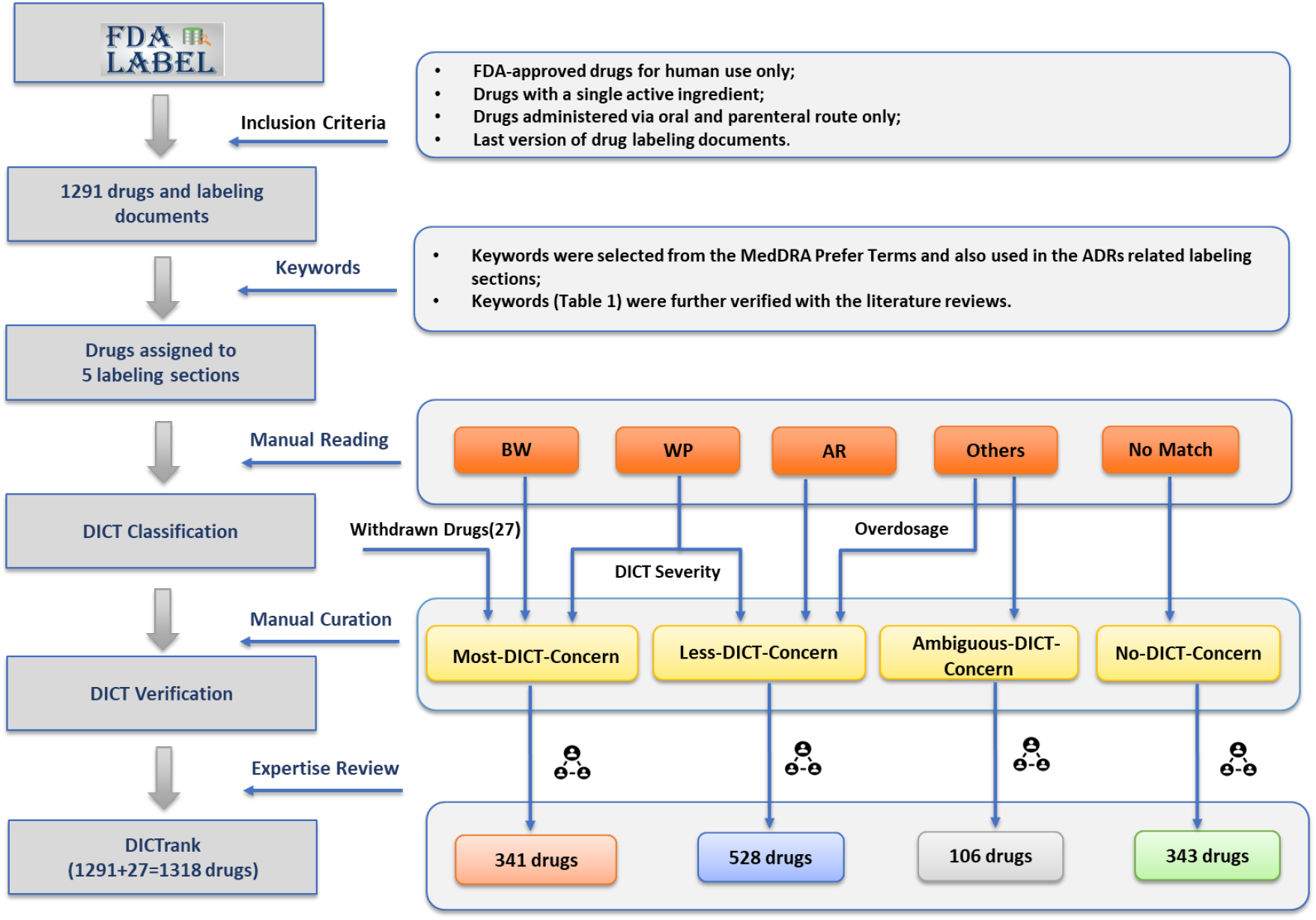
Classification scheme for drug-induced cardiotoxicity (DICT) potential on the basis of FDA-approved drug labeling documents. BW: Boxed Warning; WP: Warnings and Precautions; AR: Adverse Reactions; Others: Other labeling sections like Drug Interactions or Overdosage, and some special conditions; No-Match: none of the DICT keywords present in any ADRs related sections of drug labels. Expert Review 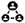: Each of the drugs was reviewed by at least two pharmacologists and the third one would be engaged in the annotation when conflicting opinions occurred.

### 1291 drugs with labeling documents were extracted

FDA-approved drug labeling is defined by the Code of Federal Regulations (21CFR201.57, www.accessdata.fda.gov/scripts/cdrh/cfdocs/cfCFR/CFRSearch.cfm?fr=201.57). Among 17 distinct sections of a drug label, ADR information are mainly documented in sections of Boxed Warning (BW), Warnings and Precautions (WP), and Adverse Reactions (AR). The same ADR described in the different sections may lead to the different clinical consideration. Specifically, the clinical implication of ADRs is in a descending order of severity [20], depending on which labeling sections are mentioned with BW > WP > AR. The BW section usually contains information of the most serious ADRs. BW section is used to concisely summarize certain contraindications or serious warnings, particularly those that can lead to death or serious injury. It is notable that multiple labeling sections may have the same ADR information, while various ADR information may occur in the same labeling section as well.

In this study, the most recent version of FDALabel [21] was used to extract labeling documents for the development of DICTrank with these criteria: (i) human prescription drugs; (ii) containing only one active ingredient; and (iii) administered orally or via parenteral route. As a result, a 1291-drug list was generated for further DICT classification.

### 1291 drugs were divided into 5 labeling sections based on DICT keywords

We developed a set of keywords (Table 1). The keywords were selected from the MedDRA Preferred Terms and also used in the ADRs related labeling sections to describe DICT. These keywords were further verified with the literature reviews [22]. Based on the keywords, the drugs were divided into 5 groups in correspondence to different labeling sections where the keyword occurs: BW, WP, AR, Others (other sections like Drug Interactions and Overdosage, and some special conditions, as show in Table 1, like patients are under cardiac disorder), and No Match (none of the DICT keywords present in any ADRs related sections of drug labels). All 1291 FDA-approved drugs were assigned to the above-mentioned five groups. When a drug hit by the keywords that present in multiple sections, the drug was assigned to the section with the following order: BW > WP > AR > Others. Full text of drug labels was subsequently reviewed manually, based on which the revision was made to these five groups by considering the context of the DICT keywords. To illustrate the revision process with an example of Dexrazoxane hydrochloride injection, in the “Warnings and Precautions” section of the labeling, it states: “Cardiac toxicity: Totect does not completely eliminate the risk of anthracycline-induced cardiac toxicity. For cardiomyopathy, monitor cardiac function before and periodically during therapy to assess left ventricular ejection fraction (LVEF). If deterioration in cardiac function occurs, consider the benefit of continued therapy against the risk of producing irreversible cardiac damage.” After thoroughly reading the entire labeling documents and considering the information provided in the “Indications and Usage” section, we placed this drug in the “Others (Special conditions)” group, even though the DICT keyword were found in the “Warnings and Precautions” labeling section.

**Table 1.**
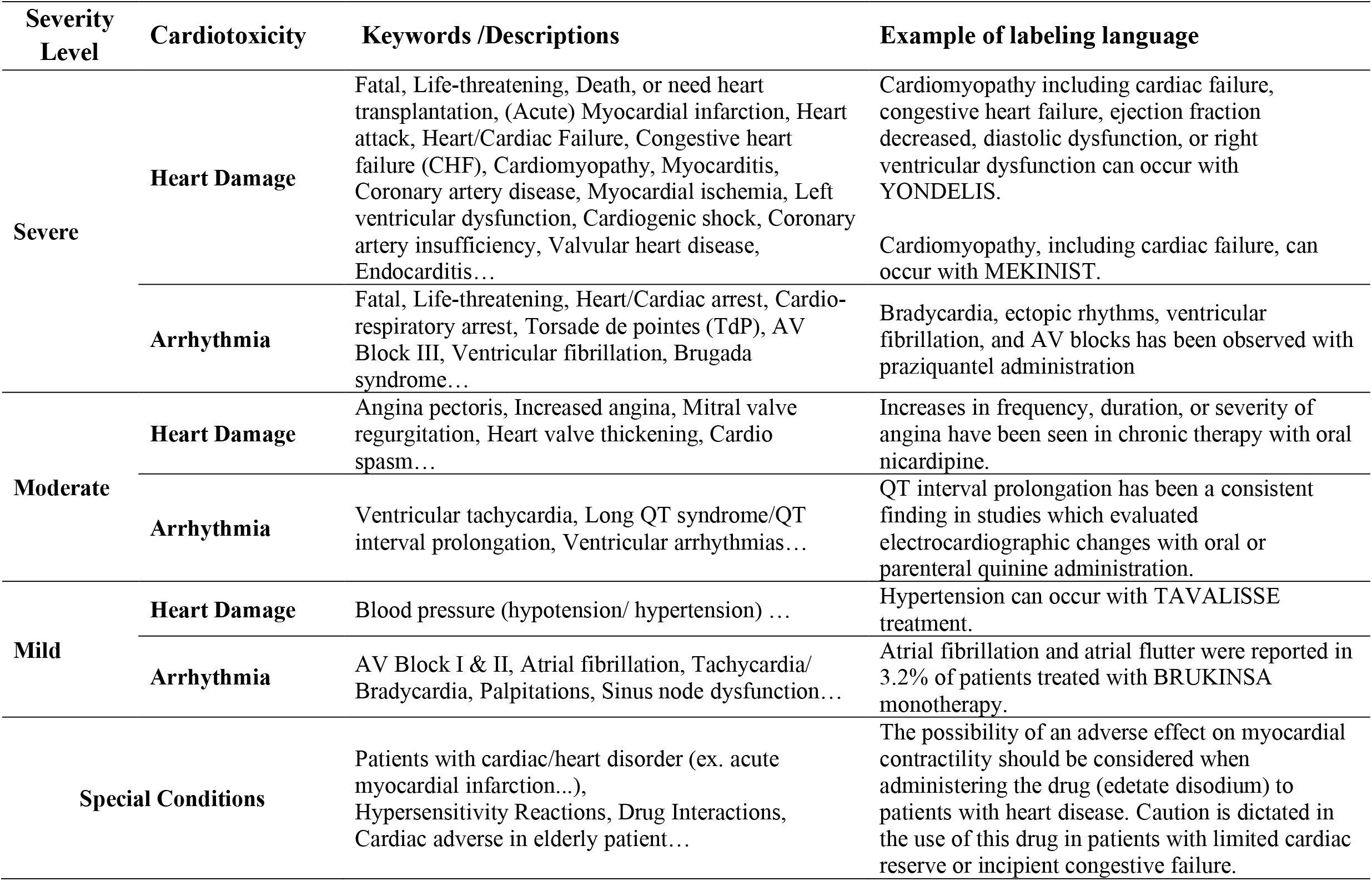
DICT categories based on the DICT description in drug labeling

### The DICT severity was considered to develop DICTrank

The description of DICT severity in the labeling language varies among drugs in the same labeling section. Considering that DICT severity has a tremendous significance in the decision making for clinical application and the review process [4,22-25], we developed a three-level system to assign all DICT keywords into three DICT severity categories: severe, moderate, and mild. As shown in Table 1, severe DICT is the highest severity level, causing serious clinical outcomes, such as cardiac death, heart failure, and myocardial ischemia. Moderate DICT causes moderate clinical DICT outcomes, such as arrhythmia and angina pectoris. Mild DICT has the less DICT severity level than moderate DICT, causing mild outcomes, such as mild arrhythmia, hypotension, hypertension.

After each individual drug was assigned to a labeling section list, the three-level severity system was applied to determine this drug’s DICT severity. As shown in Figure 2, >85.11% of the BW drugs cause Severe DICT, whereas only 34.94% WP drugs and 32.83% AR drugs belong to the Severe DICT category. The distribution of severity categories between sections confirmed that the different labeling sections record ADRs with different degrees of severity in a hierarchical way, as regulated by the Code of Federal Regulations 21CFR201.57.

**Figure 2.**
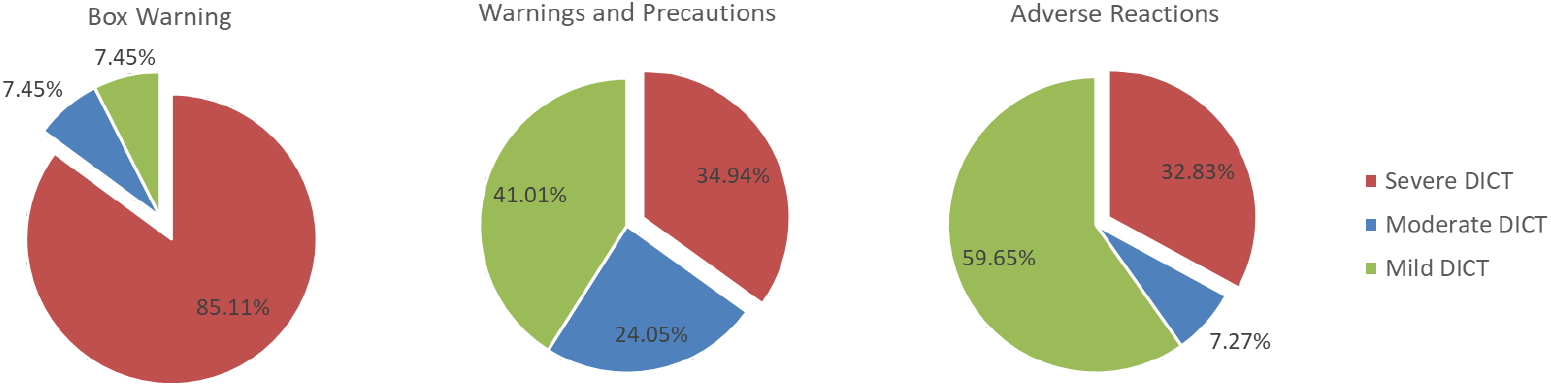
Distribution of DICT severity in the ADRs-related labeling sections

### DICTrank grouped 1318 drugs into 4 groups of DICT concern

The DICT potential was determined by integrating both labeling sections information and the severity level described above (Figure 1). A total of 1318 drugs (1291 FDA-approved drugs plus 27 withdrawn drugs) were classified into four groups depending on their DICT potentials: Most-, Less-, Ambiguous-, and No-DICT-Concern. Most-DICT-Concern group contained drugs withdrawn from the market due to severe DICT, drugs with DICT information occurring in the BW section, and WP section drugs with DICT severity in severe and moderate level. Less-DICT-Concern group contained WP section drugs which DICT severity is in mild level, drugs in AR section, and drugs in Overdosage section. Drugs with special conditions were classified as Ambiguous-DICT-Concern. Ambiguous-DICT-Concern was defined as some special conditions that there was no sufficient information in the labeling document for DICT. These special conditions (as show in Table 1) are “Patients with pre-existing cardiac/heart disorder (ex. acute myocardial infarction…),” “Hypersensitivity Reactions,” “Drug Interactions,” “Cardiac adverse in elderly patient.” For example, for “The possibility of an adverse effect on myocardial contractility should be considered when administering the drug (edetate disodium) to patients with heart disease,” we could not define the DICT risk was increased by the drug or the patients’ pre-existing cardiac disorder; therefore, the presence of DICT keywords in conjunction with these types of pre-existing conditions was considered as “Ambiguous-DICT-Concern”. Drugs with no DICT information present in any ADRs related labeling sections were grouped to No-DICT-Concern.

This classification process implicitly incorporated the factor of incidence. In clinical practice, the occurrence of severe DICT is sufficient to raise high safety concerns even though it is a rare event. When ranking a drug’s potential for DICT, the severity of DICT events in labeling was considered, with more weight given to events deemed severe as they carry greater significance [26].

To achieve the maximum rationality, expert review process was employed for the DICT verification. Each of the drugs was reviewed by at least two pharmacologists and the third one would be engaged in the annotation when conflicting opinions occurred.

The classification scheme resulted in the DICTrank drug reference list with 1318 drugs, containing 341 Most-DICT-Concern drugs, 528 Less-DICT-Concern drugs, 106 Ambiguous-DICT-Concern drugs, and 343 No-DICT-Concern drugs. A drug classified as Most-DICT-Concern was considered to have the potential to cause serious-clinical outcomes; Less-DICT-Concern denotes that a drug can cause DICT but rarely leads to serious outcomes; No-DICT-Concern denotes the associated cardiotoxicity is nonexistent; Ambiguous-DICT-Concern drugs have no sufficient information in the labeling document for cardiotoxicity.

## DICTrank analysis

The DICTrank drug list encompasses a diverse collection of therapeutic categories. The distribution of the four DICT potential levels in different therapeutic categories is illustrated in Figure 3, where the therapeutic category is defined by the second level of the World Health Organization (WHO) Anatomical Therapeutic Chemical (ATC) classification system (Table 2). The aim of the analysis was to identify drug subgroups with a higher risk of DICT. To achieve this, a Fisher exact test was performed by comparing the prevalence of Most-DICT-Concern-drugs in the investigated subgroup related to the entire DICTrank data set [17].

**Table 2.**
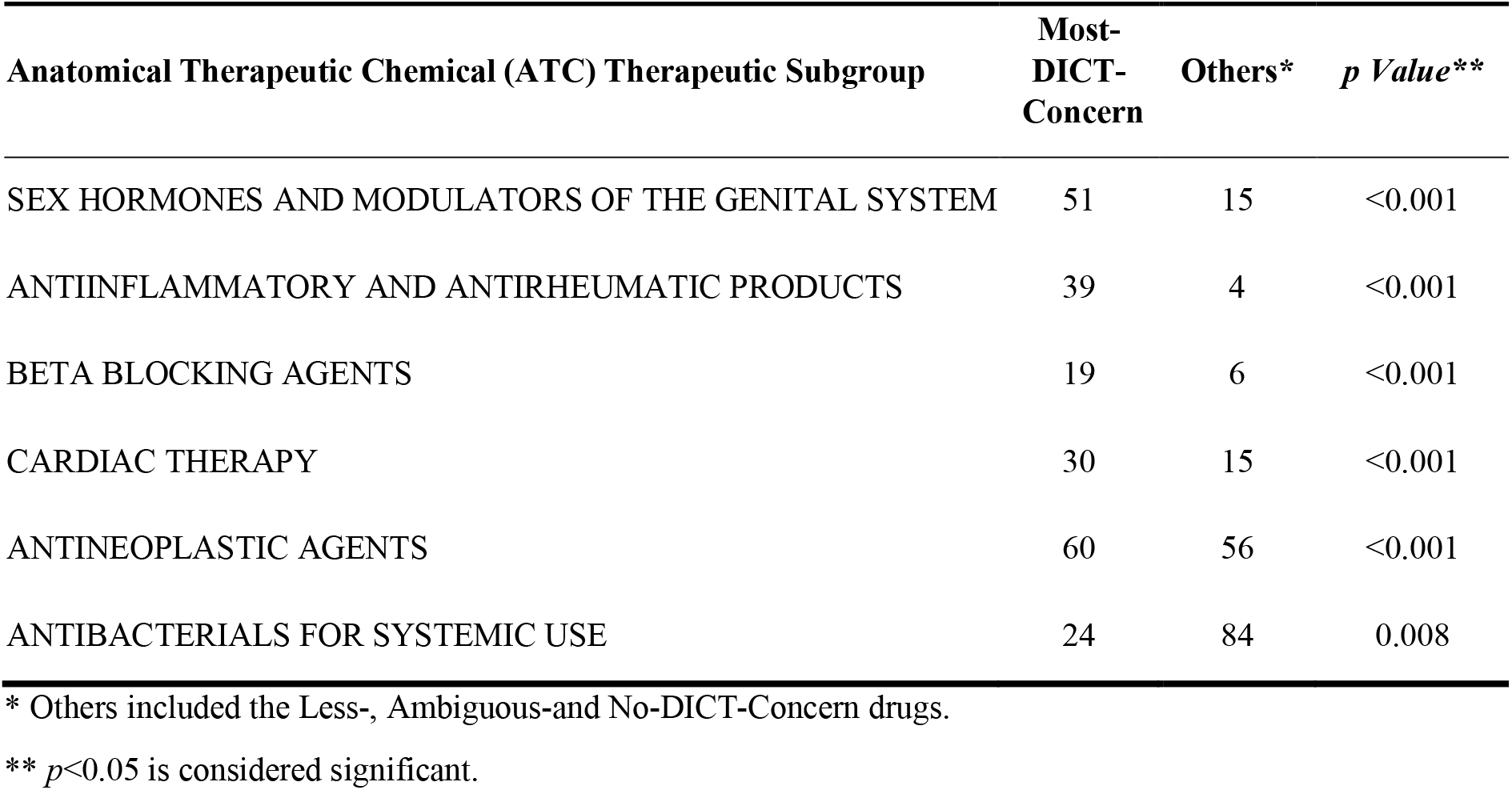
The therapeutic categories significantly enriched with the Most-DICT-Concern drugs

| Anatomical Therapeutic Chemical (ATC) Therapeutic Subgroup | Most-DICT-Concern | Others* | *p Value* ** |
| --- | --- | --- | --- |
| SEX HORMONES AND MODULATORS OF THE GENITAL SYSTEM | 51 | 15 | <0.001 |
| ANTIINFLAMMATORY AND ANTIRHEUMATIC PRODUCTS | 39 | 4 | <0.001 |
| BETA BLOCKING AGENTS | 19 | 6 | <0.001 |
| CARDIAC THERAPY | 30 | 15 | <0.001 |
| ANTINEOPLASTIC AGENTS | 60 | 56 | <0.001 |
| ANTIBACTERIALS FOR SYSTEMIC USE | 24 | 84 | 0.008 |

\* Others included the Less-, Ambiguous-and No-DICT-Concern drugs.
\*\* $p < 0.05$ is considered significant.

**Figure 3.**
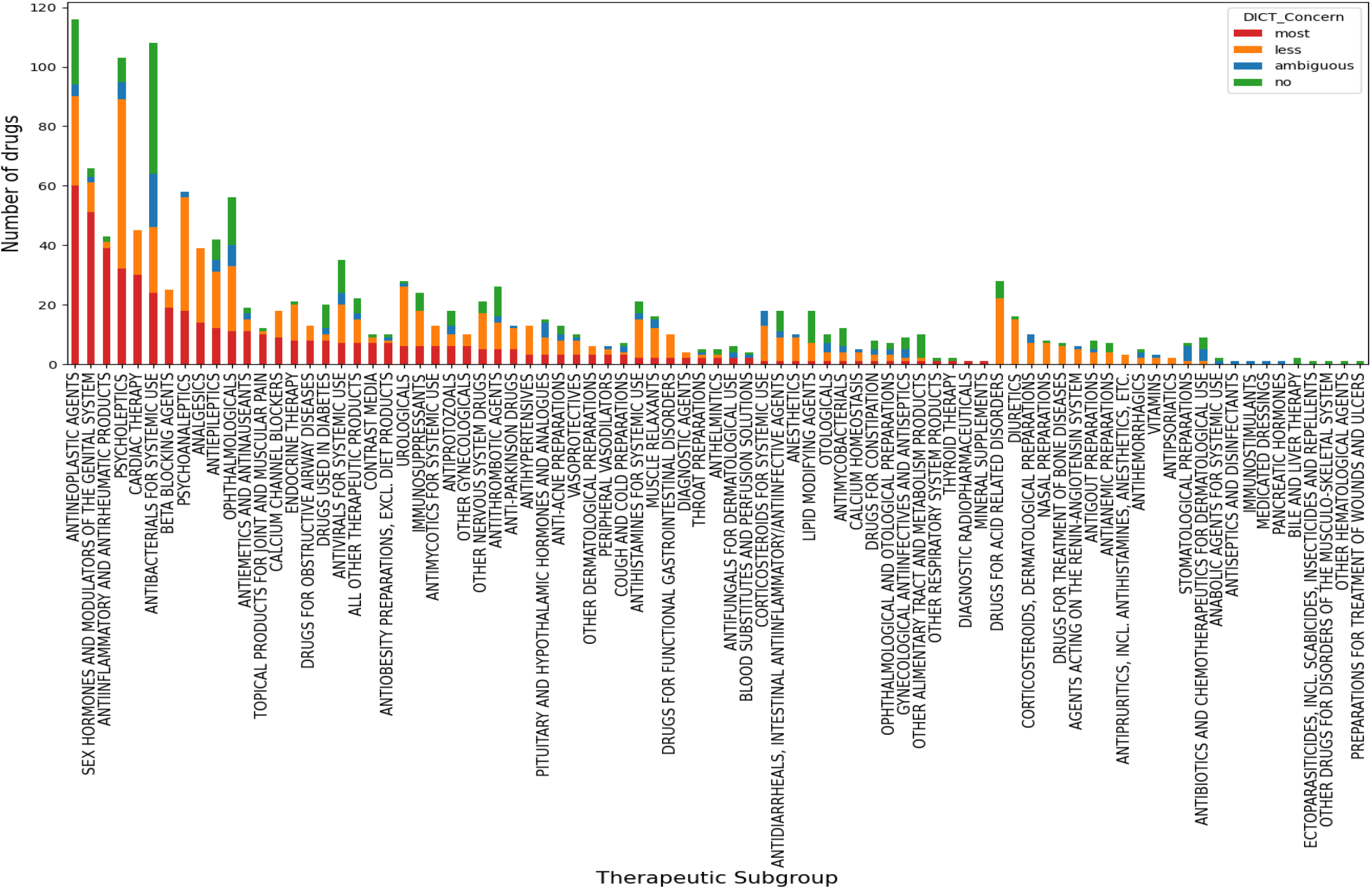
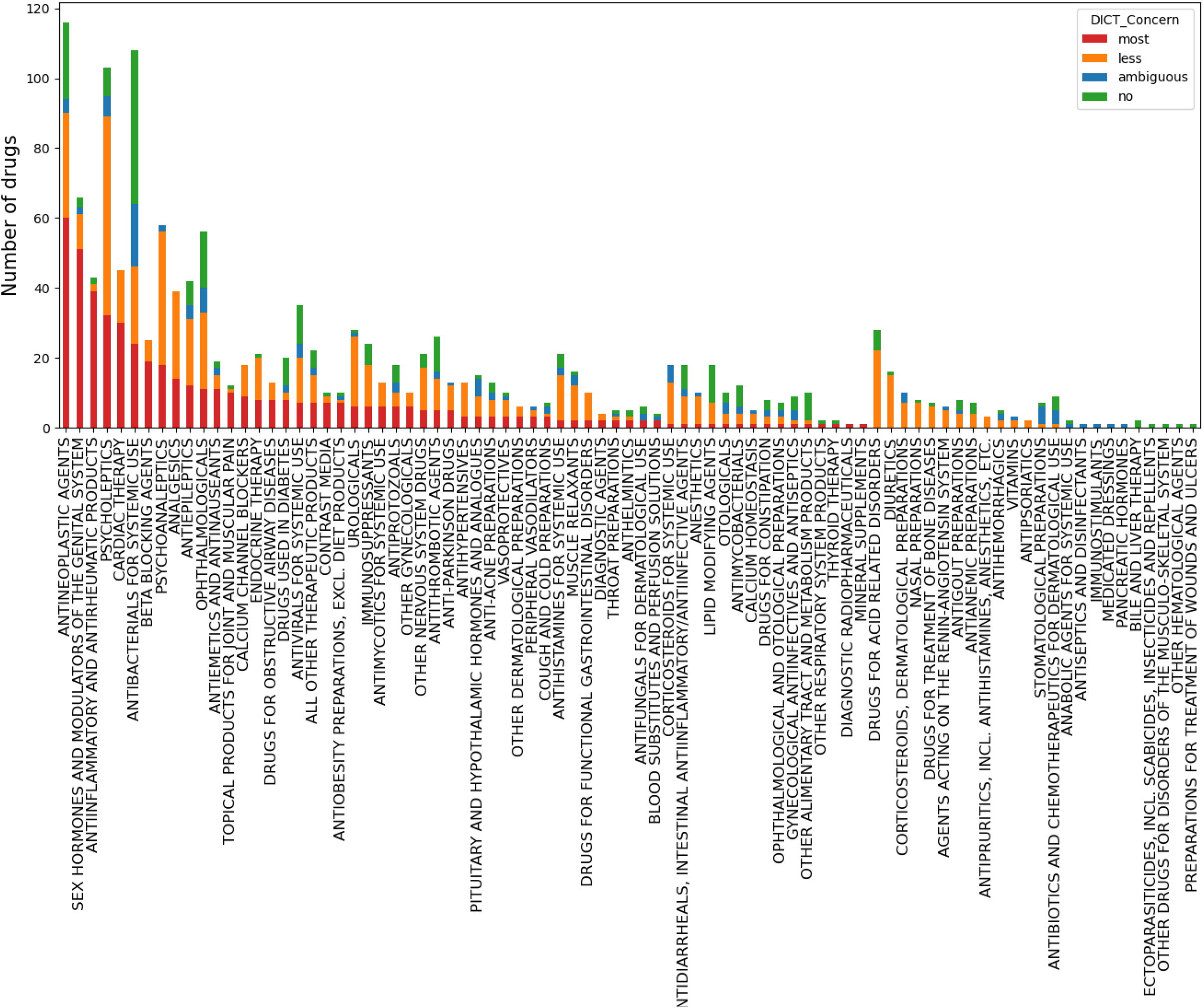
Distribution of the four DICT potential categories (Most-, Less-, Ambiguous-, and No-DICT-Concern) in different therapeutic categories

Several therapeutic categories were identified as to be significantly enriched with Most-DICT-Concern drugs. As shown in Table 2, five therapeutic categories were identified as having a significant enrichment of drugs of Most-DICT-Concern, including Antineoplastic agents, Sex hormones and modulators of the genital system, Anti-inflammatory and antirheumatic products, Beta blocking agents and Cardiac therapy. It is notable that these therapeutic categories are commonly linked with cardiotoxicity, and the profile attained aligns with findings documented in literature [4,13,27-30].

The adjustment of dosage regimens for specific subpopulations is a critical aspect of precision medicine practice, aiming to mitigate the occurrence of unforeseen ADRs [31]. Overdosing has been widely reported to be one of the risk factors for DICT [32,33]. We conducted a systematic investigation of dosage distribution across different therapeutic categories, where the drug daily dose (DDD) information was extracted from ATC/DDD Index 2023 (www.whocc.no/atc_ddd_index/) [14,21]. As shown in Figure 4, the Most-DICT-Concern drugs in some therapeutic categories (i.e., J01 – Antibacterials for systemic use, M01 – Anti-inflammatory and antirheumatic products, C07 – Beta blocking agents, L01 – Antineoplastic agents, C01 – Cardiac therapy) have relatively higher daily doses, suggesting more attention should be paid to the dosage when using these drugs in clinical practice.

**Figure 4.**
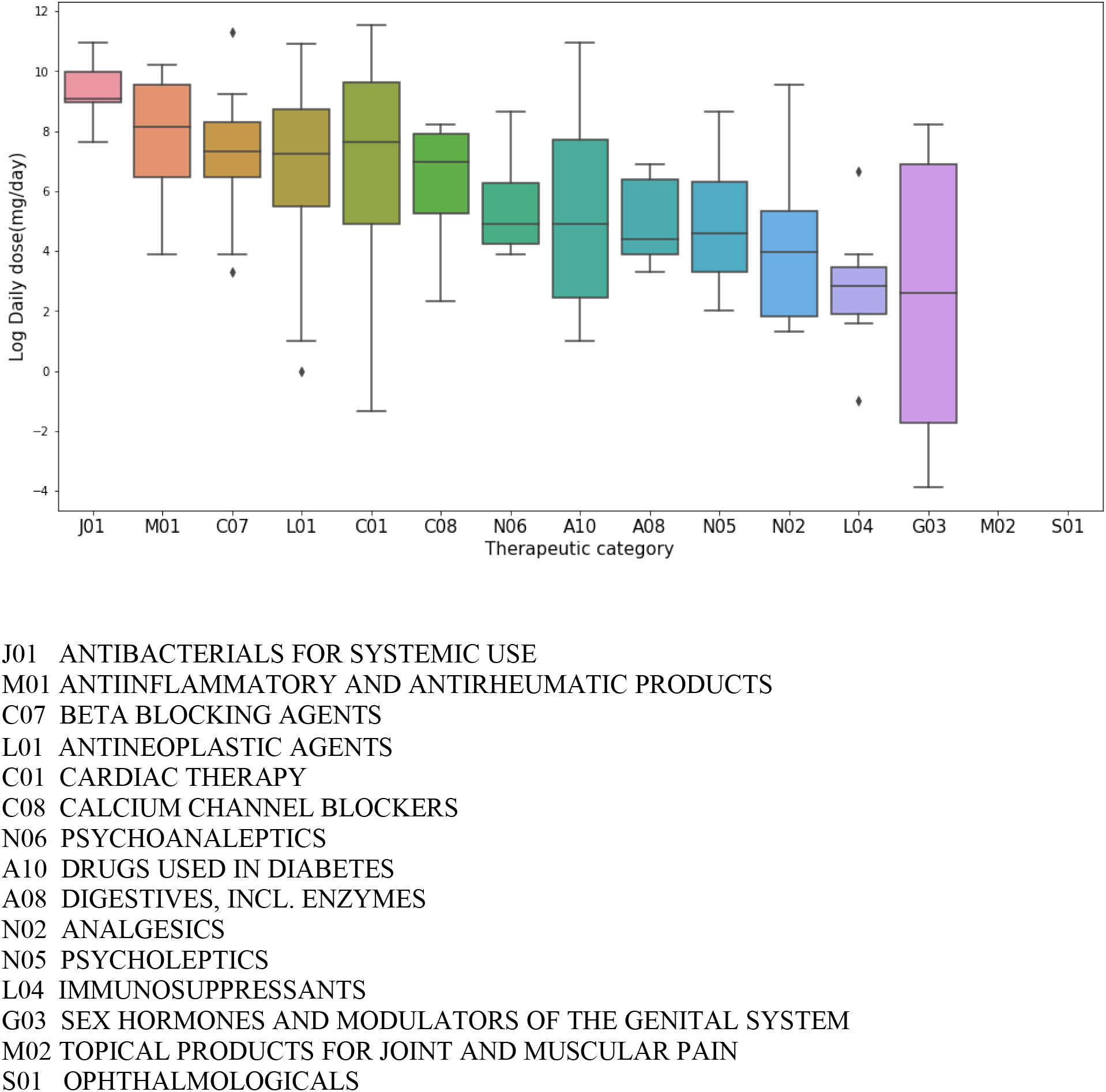
Human daily dose distribution of Most-DICT-Concern drugs based on WHO ATC/DDD Index 2023. The second level of ATC Classification System was used to represent the therapeutic categories of Most-DICT-concern drugs. The corresponding human daily dose was collected for Most-DICT-concern drugs used in multiple therapeutic categories. The daily dose distribution of thirteen therapeutic categories was then box plotted.

## Discussion

DICT is a significant safety concern in both drug safety and clinical diagnosis. It ranks among the most frequently reported serious ADRs during clinical practice and drug development [35]. For instance, comprehensive reviews that cover various durations and regions have indicated that DICT is a leading cause of drug withdrawals from the market [3, 36, 37]. Most of studies have been focused on specific DICT adverse events or mechanisms such as QT prolongation and hERG-related cardiotoxicity, where several drug lists have been reported [5,14,34]. However, to the best of our knowledge, there have been no reports aggregating these datasets to provide an overall assessment of the potential for DICT with a large number of drugs and across a broad range of drug classes.

In this study, we developed the DICTrank dataset, which is the largest and most comprehensive reference drug list for DICT safety assessment. We utilized a systematic process that involved keyword searches and manual curation of FDA labeling documents to create this dataset. Our two-step approach assessed a drug’s potential for DICT by considering the presence of DICT keywords in the labeling section and the severity of observed DICT for the drug. As a result, we annotated 1318 drugs in the DICTrank dataset and classified them into four ranks: Most-DICT-Concern (341 drugs), Less-DICT-Concern (528 drugs), Ambiguous-DICT-Concern (106 drugs), and No-DICT-Concern (343 drugs).

There has been concerns to utilize the labeling documents for safety assessment[16], which has been debated in our previous study on drug-induced liver injury [16,17]. For instance, there is a concern that adverse drug reaction keywords in labeling documents may have evolved over time. However, many experts believe that labeling documents still represent the consensus of experts at the time of their creation, reflecting the serious safety concerns of regulators and drug manufacturers [16]. It is ‘the closest that one can get to the truth regarding the scientific information known about a drug’ [19]. It is worthwhile to mention that DICT potential in this study is a matter of clinical concern rather than a regulatory decision [21]. The current version of DICTrank comprises FDA-approved drug labeling data (1291 drugs) and includes known withdrawn drugs (27 drugs) due to DICT. The dataset can be expanded in the future by incorporating labeling data from other countries.

It is worth highlighting that DICTrank focuses specifically on drug-induced cardiotoxicity (DICT), which refers to adverse effects or damage to the heart caused by drugs. It centers on the direct toxic effects that affect the structure or function of the heart. In contrast, cardiovascular toxicity has a broader scope and encompasses adverse effects or toxicity that affect the entire cardiovascular system, including the heart, blood vessels, and blood circulation. It considers the toxic effects on the cardiovascular system as a whole, rather than solely targeting the heart. Our focus on DICT addresses the need for assessing and managing potential cardiac toxic effects in therapeutic scenarios where heart injury is a primary concern, such as certain medications, targeted therapies, and radiation therapies.

Considering the complex molecular process of DICT, the underlying mechanism of DICT has yet to be fully understood. A high-quality reference drug list containing comprehensive DICT information is crucial for better studying underlying mechanisms, discovering DICT biomarkers, and developing effective diagnosis and prognosis strategies. In the current era of big data, alternative approaches to using vertebrate animals for drug safety assessment have gained increasing importance. Accurate and consistent annotation of drugs is essential to develop new methodologies utilizing advanced technologies, such as high-throughput, high-content, organ-on-a-chip, 3D cellular models, stem cell-derived cardiomyocyte models, omics, quantitative structure-activity relationships (QSARs), as well as data-driven and mechanism-driven modeling [35-39].

To the best of our knowledge, the DICTrank dataset provided in this study is currently the largest publicly available annotated dataset for DICT. It has the potential to assist in the development of predictive models for early detection of DICT risk in drug development, as well as in clinical and epidemiological investigations.

## Disclaimer

The opinions expressed in this paper are those of the authors, and do not necessarily reflect the position of U.S. Food and Drug Administration.

The dataset is currently under review and may be provided upon request.

## Acknowledgements

This research is part of the SafetAI initiative supported by FDA’s Center for Drug Evaluation and Research (CDER).

YQ is grateful for the support by an appointment to the Research Participation Program at the National Center for Toxicological Research (NCTR) of the U.S. FDA through the Oak Ridge Institute for Science and Education (ORISE).

We thank Dr. Feng Wei (Cardiologist, MD, PhD) from the Department of Structural Heart Disease, the First Affiliated Hospital of Xi’an Jiaotong University (China) for consulting on DICT classification.

We thank Dr. Jürgen Borlak from the Center of Pharmacology and Toxicology, Hannover Medical School (Germany) for inspiring discussion during the early stage of this project.

